# Electrostatics and Local Aromatic Residues Govern Lipid Binding and Membrane Penetration of Synaptotagmin C2 Domains

**DOI:** 10.64898/2026.07.09.737582

**Authors:** Dong An, Manfred Lindau

## Abstract

Synaptotagmins (Syts) are Ca²⁺-sensing exocytosis regulators whose tandem C2 domains interact with phosphoinositides and membranes to trigger neurotransmitter and hormone release. Although Ca²⁺ binding is known to enhance C2 domain-membrane interactions, the sequence determinants governing lipid binding and membrane penetration across Syt isoforms remain incompletely understood.

Here, we performed MARTINI coarse-grained molecular dynamics simulations of isolated C2A and C2B domains from eight Ca²⁺-sensing Syt isoforms (Syt1, Syt2, Syt3, Syt5, Syt6, Syt7, Syt9, and Syt10) interacting with phosphatidylinositol 4,5-bisphosphate (PIP₂)-containing plasma membranes. To systematically modulate electrostatic properties, we introduced partial and full charge-flip mutations at conserved acidic residues within the calcium-binding loops (CBLs). By integrating simulations across multiple isoforms and charge states, we sought to identify the dominant sequence determinants governing membrane interactions.

We found that PIP₂ binding to both, CBLs and polybasic patches (PBs), is associated with loop net charge, yielding correlations > 0.95 across all isoforms. However, membrane penetration is not sufficiently explained by loop net charge alone. The local phenylalanines additionally increase membrane penetration independent of loop net charge.

Together, these findings establish a comprehensive electrostatic–aromatic framework where loop net charge governs PIP₂ binding, whereas loop net charge and local phenylalanine enrichment jointly govern membrane penetration across Syt C2 domains.

## Introduction

C2 domains are membrane-binding modules found in numerous eukaryotic signaling proteins, including protein kinase C (PKC), Synaptotagmins (Syts), mammalian uncoordinated-13 protein (Munc13), Rab interacting molecules (RIMs), and Extended Synaptotagmins (E-Syts) [1–4]. These proteins mediate diverse signaling processes through membrane interactions, often coupling lipid binding to intracellular Ca²⁺ signaling.

One prominent example of C2-domain function is exocytosis, including neurotransmitter release and hormone secretion, through membrane fusion mediated by soluble N-ethylmaleimide–sensitive factor attachment protein receptor (SNARE) complexes [5,6]. This process is tightly regulated by C2 domain containing proteins such as RIMs, Munc13 and Syts from SNARE complex assembly, vesicle docking, priming, to membrane fusion evoked by Ca²⁺ signals, which are sensed by Syt C2 domains [7,8].

To date, 17 synaptotagmin isoforms have been identified in the *Homo sapiens* and *Mus musculus* genomes [4]. Among these, several isoforms function as Ca²⁺ sensors, including Syt1, Syt2, Syt3, Syt5, Syt6, Syt7, Syt9, and Syt10 with different Ca²⁺-sensing kinetics responses to Ca²⁺ influx [9,10]. This classification is functionally relevant, as distinct Syt isoforms are selectively enriched in specific subsets of secretory vesicles and dense-core granules [11]. For example, Syt1 and Syt2 primarily mediate synchronous Ca²⁺-evoked neurotransmitter release [12], whereas Syt3 and Syt7 have been implicated in asynchronous release pathways [13].

Despite extensive experimental characterization of Syts, the sequence determinants governing lipid binding and membrane penetration of C2 domains remain poorly understood. This gap is largely attributable to the limited spatiotemporal resolution of current experimental approaches [6]. Even among fast Ca²⁺ sensors, Syt1 and Syt2 display distinct Ca²⁺ sensitivities and release kinetics, underscoring the presence of isoform-specific regulatory mechanisms [12].

Recent experimental studies have begun to elucidate the structural and biophysical basis of Syt function. NMR experiments from the Rizo group demonstrated that Ca²⁺ binding to the C2B domain of Syt1 enhances its interaction with anionic lipids, particularly PIP₂ [14]. Building on this data, a mechanistic model was proposed in which Syt1, bound to the SNARE complex at the primary interface [15], partially dissociates upon Ca²⁺-induced C2B loop penetration into the membrane, thereby levering the SNARE complex to promote membrane fusion [16]. Complementary work from the Chapman group showed that Ca²⁺-bound C2B loops of Syt7 penetrate more deeply into anionic membranes and modulate lipid order, highlighting isoform-specific membrane interactions [17].

Nevertheless, several critical questions remain unresolved. Which sequence features govern lipid binding and membrane penetration by C2 domains? How do these features contribute to the distinct membrane-interaction properties of C2A and C2B domains? What is the potential division of labor between C2A and C2B during membrane engagement?

To address these questions, we performed MARTINI coarse-grained molecular dynamics simulations of isolated C2A and C2B domains from Syt1, Syt2, Syt3, Syt5, Syt6, Syt7, Syt9, and Syt10 interacting with PIP₂-containing plasma membranes (Fig. 1). To systematically modulate the electrostatic properties of calcium-binding loops (CBLs), we introduced partial and full charge-flip mutations at conserved acidic residues. We quantified the average number of PIP₂ molecules bound to the CBLs and PBs, as well as the membrane penetration depth, by averaging these quantities over the final 1 μs of each trajectory. By integrating simulations across multiple isoforms and charge states, we identified the dominant sequence determinants governing these properties.

**Figure 1.**
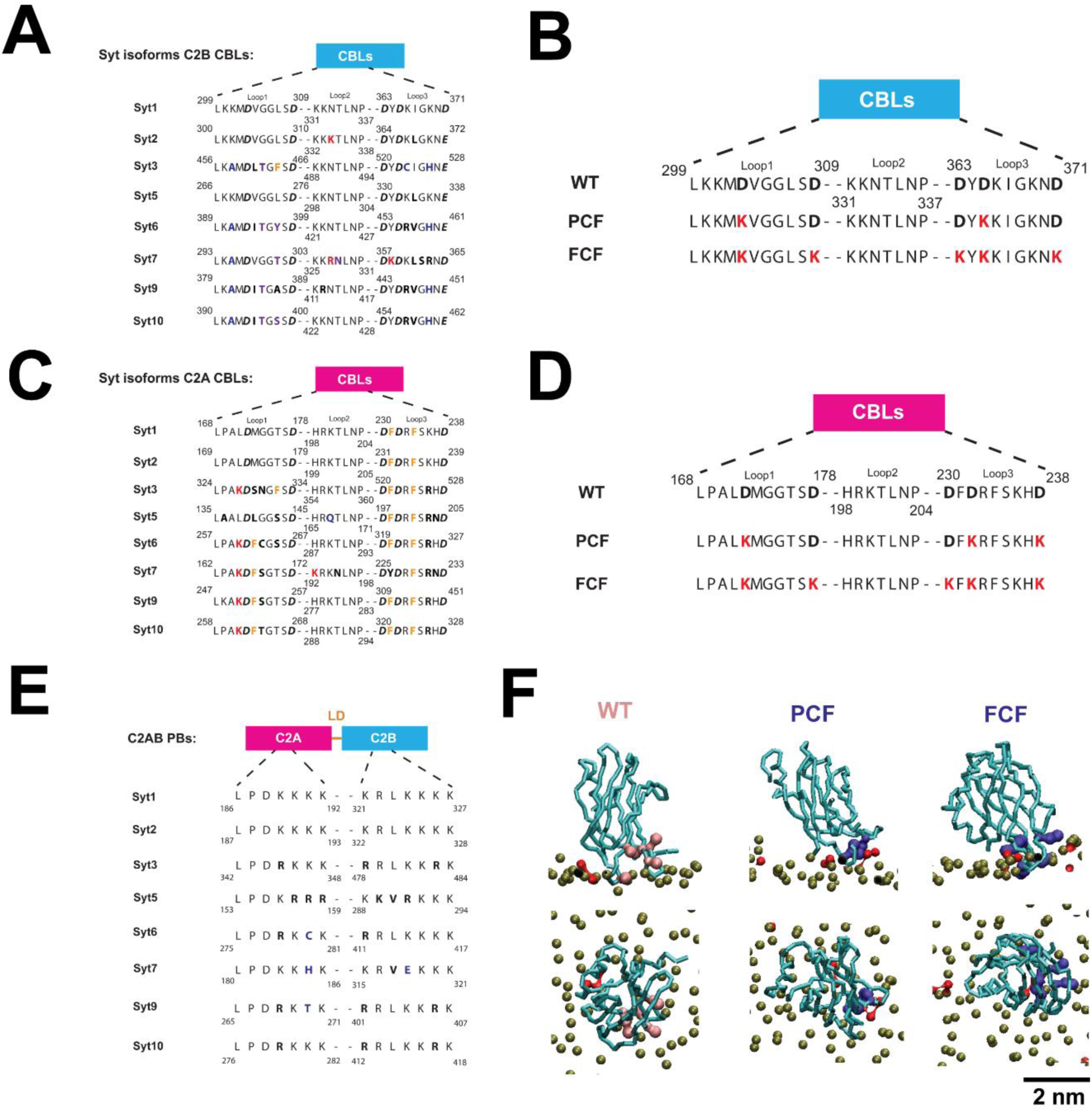
Synaptotagmin (Syt) C2AB sequences, charge-flip mutations, and simulation setup. **(A)** Multiple sequence alignment of Ca²⁺-binding loops (CBLs) in the C2B domains of Syt1, Syt2, Syt3, Syt5, Syt6, Syt7, Syt9, and Syt10. Residue substitutions are color-coded as follows: gain of positive charge (red), loss of positive charge (blue), substitutions preserving hydrophobic or charge character (black bold), and hydrophobic-to-hydrophilic substitutions (purple). Conserved Ca²⁺-coordinating Asp/Glu residues are shown in bold italics. Phe residues are colored orange. **(B)** Schematic representation of charge-flip mutations introduced into Syt1 C2B CBLs. Partial and full charge-flip mutations (PCF and FCF) correspond to DK2 and DK5, respectively. Equivalent mutations were introduced into corresponding residues of other Syt isoforms. **(C)** Multiple sequence alignment of CBLs in the C2A domains, formatted as in **(A)**. Compared with C2B domains, C2A domains contain more aromatic residues, particularly Phe. **(D)** Schematic representation of charge-flip mutations introduced into Syt1 C2A CBLs. PCF and FCF correspond to DK3 and DK5, respectively. Equivalent mutations were introduced into corresponding residues of other isoforms. **(E)** Multiple sequence alignment of PBs in the C2A and C2B domains of the indicated Syt isoforms. **(F)** Representative initial configurations of Syt1 C2B–plasma membrane (PM) simulations in the Ca²⁺-free (WT), PCF, and FCF states. C2B is shown in cyan, phospholipid phosphate beads in brown, PIP₂ phosphate beads in red, conserved Asp/Glu residues in pink, and charge-flipped Lys residues in violet. Loops 1 and 3 were initially positioned in contact with the membrane to facilitate sampling of membrane-bound conformations. Equivalent setups were used for all C2A and C2B simulations.

Our analyses reveal that PIP₂ binding is primarily determined by loop net charge. In contrast, membrane penetration cannot be explained by electrostatics alone. Instead, membrane penetration is jointly governed by loop net charge and local phenylalanine enrichment. Together, these findings establish a comprehensive quantitative electrostatic–aromatic framework linking C2-domain sequence composition to membrane interactions across Syt isoforms.

## Results

### PIP₂ binding to CBLs and PBs is associated with loop net charge

Previous studies have shown that PIP₂ interacts with both the CBLs and PBs of Syt C2 domains through Ca²⁺-dependent and Ca²⁺-independent mechanisms, respectively [14,18]. Because charge-flip mutations selectively increase the net charge of CBLs mimicking Ca^2+^ binding while leaving PB charge unchanged, we first examined whether net charge influences PIP₂ binding.

We quantified the average numbers of PIP₂ molecules bound to CBLs and PBs for the C2 domains of Syt1, Syt2, Syt3, Syt5, Syt6, Syt7, Syt9, and Syt10 under WT, partial charge-flip (PCF), and full charge-flip (FCF) conditions over the last 1 μs (Fig. S1-3, Table S1). To show that the PIP₂ binding is stable over the last 1 μs (Fig. 2A, S1), we also analyzed separately the time intervals of 2.0–2.5 μs and 1.5–2.0 μs (Fig. S2), and no obvious differences were observed. As representative examples, Syt1 and Syt7, which are widely studied as fast and slow Ca²⁺ sensors, respectively, both exhibited increased CBL–PIP₂ interactions following PCF and FCF mutations (Fig. 2B). Similar trends were observed across most other isoforms (Fig. S2A-C), although the magnitude of the increase varied among isoforms.

**Figure 2.**
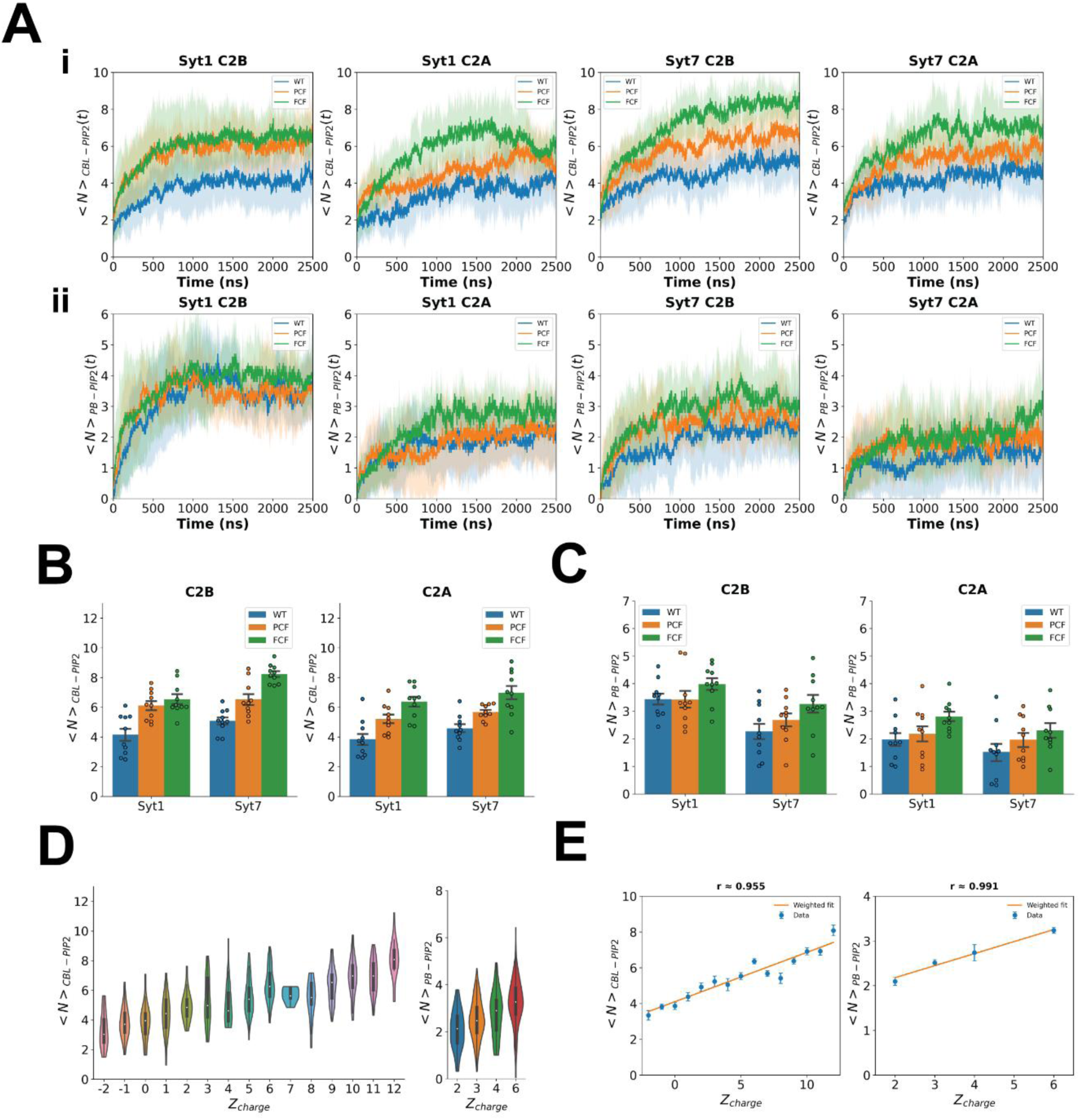
Loop net charge is the dominant determinant of PIP₂ binding. **(A)** Representative trajectories of CBL–PIP₂ **(i)** and PB-PIP2 **(ii)** binding for Syt1 and Syt7 C2A and C2B domains under WT, PCF, and FCF conditions. Solid lines represent the mean values from *n* = 10 independent simulations, and shaded regions indicate SDs. Average trajectories for other Syt C2 domains are shown in Fig. S1. **(B)** Average numbers of PIP₂ molecules bound to CBLs in the C2B and C2A domains of Syt1 and Syt7 during 1.5–2.5 μs. Bars represent mean ± SEM and dots correspond to ten independent simulations. Numbers in parentheses indicate CBL net charges at WT state. Corresponding analyses for other Syt isoforms are shown in Fig. S2. Individual details are summarized in Table S2. **(C)** Average numbers of PIP₂ molecules bound to PBs in the same simulations during 1.5–2.5 μs. Bars represent mean ± SEM and dots correspond to ten independent simulations. Numbers in parentheses indicate PB net charges. Corresponding analyses for other Syt isoforms are shown in Fig. S3. Individual details are summarized in Table S2. **(D)** Distributions of average PIP2 molecules bound to calcium-binding CBLs (left) and polybasic patches (PBs, right) over 1.5–2.5 μs regrouped according to the primary sequence descriptor (Figure 1A-E), namely the net charge (*Z_c_*_ℎ*arge*_ ) of the CBLs or PBs. Similar regrouping analyses using the averages from 1.5–2.0 μs and 2.0–2.5 μs are shown in Fig. S4A. Sample sizes and statistics for each charge group are provided in Table S3 for CBLs and Table S4 for PBs. **(E)** Linear regressions between the mean number of bound PIP2 molecules versus the net charges of CBLs (left) or PBs (right). Data from all C2 domains and charge states were regrouped according to net charge, as shown in **(D)**, before regression analysis. Blue dots represent the mean numbers of bound PIP₂ averaged across all C2 domains sharing the same net charge. Error bars indicate SEM. Orange lines indicate the best-fit linear regression.

In contrast, PIP₂ interactions with PBs were largely unaffected by PCF and FCF mutations (Fig. 2C and Fig. S3A-C). This result is expected because PB net charge remains unchanged in all charge-flip variants, despite the close spatial proximity between PBs and the CBLs by 6-7 amino acids (Fig. 1A, C, E).

Together, these observations suggest that PIP₂ interactions are primarily associated with the net charge of the protein region involved.

### Loop net charge is the major determinant of PIP₂ binding to CBLs and PBs

To identify the dominant determinants of PIP₂ binding, we regrouped all simulations according to the net charges of the corresponding protein regions, irrespective of C2 domain type, isoform identity, or charge-flip state.

For CBLs, the 480 simulations were regrouped into 15 charge groups spanning net charges from −2 to 12 (Figure 2D left, S4A, Table S3). The mean numbers of PIP₂ molecules bound to CBLs exhibited a strong positive correlation with CBL net charge (Fig. 2E left). Pearson correlations were approximately 0.96, indicating that net charge accounts for the variation in CBL–PIP₂ interactions. Similar results were obtained using the partial time windows of 2.0–2.5 μs and 1.5–2.0 μs (Fig. S4B).

Similarly, simulations were regrouped into four PB charge groups with net charges of 2, 3, 4, and 6 (Figure 2D right, Table S4). The mean numbers of PIP₂ molecules bound to PBs were even more strongly associated with PB net charge, yielding Pearson correlations of approximately 0.99 in both equilibrium windows (Fig. 2E right, S4B). Similar results were obtained using the partial time windows of 2.0–2.5 μs and 1.5–2.0 μs (Fig. S4B).

To evaluate whether these relationships were also robust to ranking rather than just absolute values, we additionally calculated Spearman correlations. Spearman and Pearson correlations were nearly identical for both CBLs and PBs (Fig. S4C–D), indicating that the observed charge dependence is both monotonic and approximately linear. Furthermore, leave-one-out analyses yielded highly similar correlations (Fig. S5), demonstrating that these relationships were not driven by any individual charge group.

Together, these results identify loop net charge as the dominant first-order determinant of PIP₂ interactions in both CBLs and PBs.

### Net charge alone does not fully explain variation in CBL membrane penetration

The second property examined in this study was CBL membrane penetration depth, defined as the vertical distance between the vertical position of the IC leaflet head group plane (*Z_ref_*) and the minimum z-coordinate among all CBL beads (*Z^min^*) (Fig. 3A) as

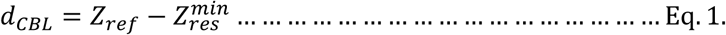

**Figure 3.**
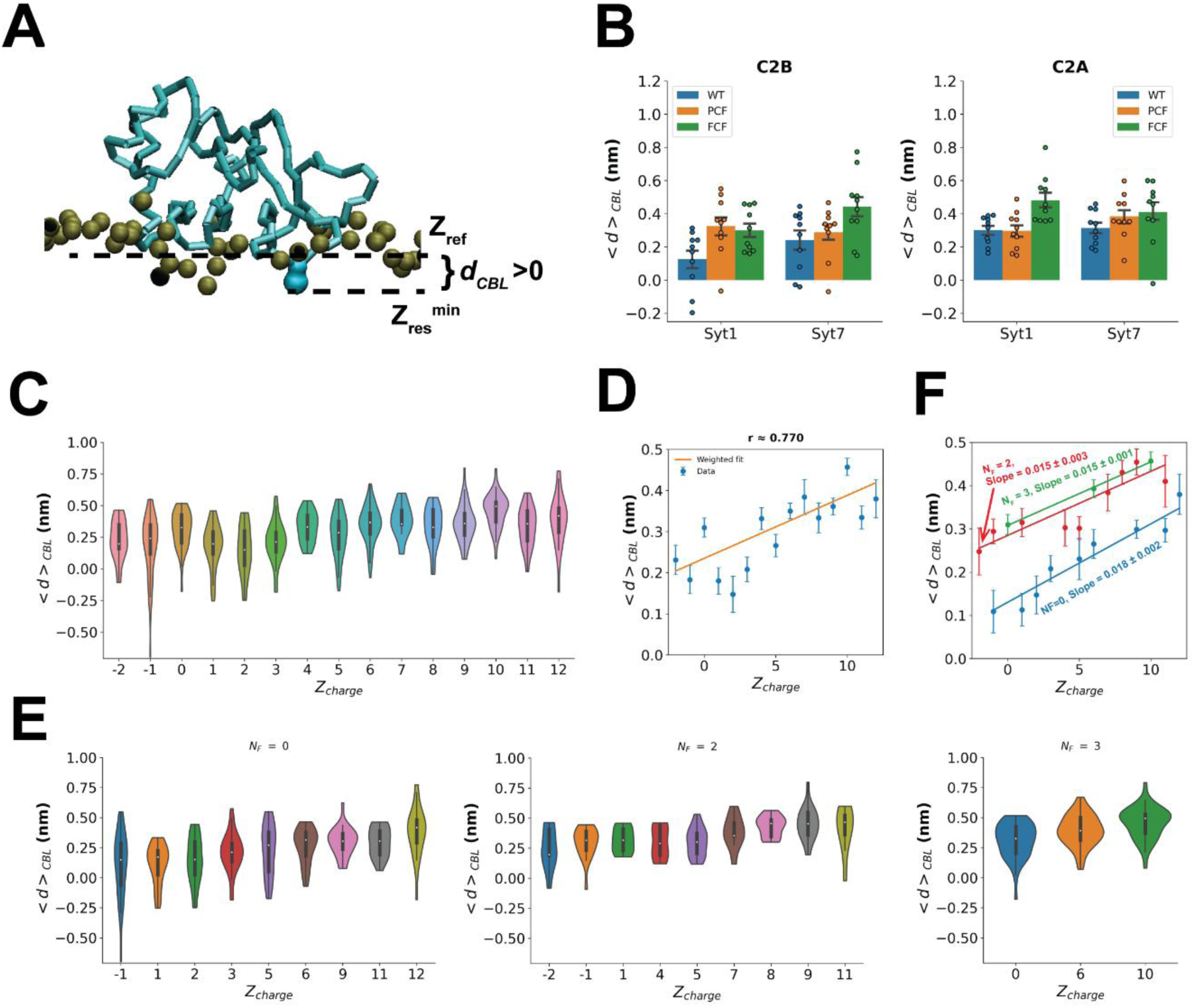
Loop net charge and local phenylalanine enrichment jointly govern membrane penetration. **(A)** Definition of CBL membrane penetration depth. A representative snapshot of the Syt1 PCF C2B domain bound to the plasma membrane is shown. The penetration depth (*d_CBL_*) was defined as the difference between the average *z*-coordinate of the lipid headgroup plane (*Z_ref_*) and the minimum *z*-coordinate among all CBL beads (*Z^min^*) (Eq. 1). Positive values indicate insertion of CBL residues into the membrane hydrophobic region. **(B)** Average CBL penetration depths in the C2B and C2A domains of Syt1 and Syt7 during 1.5–2.5 μs. Values represent mean ± SEM from *n* = 10 independent simulations. Numbers in parentheses indicate the net charges of the CBLs in the WT state. CBL penetration depths of all Syt isoforms are shown in Fig. S7. Individual details are summarized in Table S5. **(C)** Average CBL penetration depths were regrouped according to CBL net charge (*Z_c_*_ℎ*arge*_ ), irrespective of C2 domain type or Syt isoform. Sample distributions are shown for the final 1 μs (1.5–2.5 μs). Although membrane penetration generally increased with increasing loop net charge, substantial overlap remained among charge groups, suggesting that additional sequence determinants contribute to membrane penetration. **(D)** Linear regression between the mean CBL penetration depth and CBL net charge after regrouping according to *Z_c_*_ℎ*arge*_ . Blue dots represent the mean penetration depths averaged across all C2 domains sharing the same net charge, and error bars indicate SEM. The orange line represents the best-fit linear regression. Loop net charge alone explained approximately 55% of the variation in membrane penetration. **(E)** membrane penetration distributions after stratification according to the number of local phenylalanine residues (*N_F_* = 0, 2, or 3). Within each *N_F_* subgroup, data were further regrouped according to CBL net charge. Separating simulations according to local phenylalanine enrichment substantially reduced the overlap among charge groups, indicating that local aromatic composition accounts for much of the residual variation not explained by loop net charge alone. **(F)** Linear regressions between the mean CBL penetration depth and CBL net charge within each *N_F_* subgroup. Similar slopes (mean ± SE) were obtained for all three *N_F_* categories, whereas increasing *N_F_* primarily shifted the intercept toward larger membrane penetration depths. These results demonstrate that loop net charge remains the dominant determinant of membrane penetration within each local phenylalanine category, while local phenylalanine enrichment provides an additional independent contribution, together forming a simple two-feature model for predicting membrane penetration. Slopes are in the unit of nm/*Z_c_*_ℎ*arge*_ .

Positive values indicate insertion of CBL residues into the membrane hydrophobic region, whereas negative values indicate that CBL residues remain above the membrane interior.

We quantified the average penetration depths of CBLs from the C2 domains of Syt1, Syt2, Syt3, Syt5, Syt6, Syt7, Syt9, and Syt10 under WT, partial charge-flip (PCF), and full charge-flip (FCF) conditions (Fig. S6-7 and Table S4). No obvious differences were observed over the last 1 μs in the average trajectories (Fig. S6). Representative examples from Syt1 and Syt7 showed that increasing CBL charge generally increased membrane penetration in the last 1 μs equilibrium windows (Fig. 3B), although the magnitude of the changes varied among C2 domains and isoforms (Fig. S6-7).

To identify the dominant determinants of membrane penetration, all 480 simulations were regrouped according to CBL net charge, yielding 15 charge groups spanning values from −2 to 12 (Fig. 3C, Table S2). Mean penetration depths exhibited a positive dependence on CBL net charge in both equilibrium windows (Fig. 3D). However, the resulting Pearson correlations were only 0.77 for the last 1μs interval. Thus, unlike PIP₂ binding (Fig. 2C-D, S3), net charge alone can only yield R^2^ of 0.57, meaning that only 57% of the changes of membrane penetration can be explained by net charge alone. Similar results were yielded for 1.5–2.0 μs and 1.5–2.0 μs intervals (Fig. S8A-B).

These observations suggest that although electrostatic interactions contribute substantially to membrane penetration, additional sequence-derived factors must also participate in determining CBL membrane interactions.

### At the same numbers of local Phenylalanine residues, net charge becomes the major determinant of CBL membrane penetration

Comparison of the C2A and C2B sequence alignments revealed a striking difference in the local environments surrounding the conserved acidic residues of the calcium-binding loops (Fig. 1A, C). Whereas C2B domains are enriched in aliphatic residues such as Leu, Ile, and Val, C2A domains frequently contain one or two phenylalanine residues adjacent to conserved Asp or Glu residues. Because aromatic substitutions generally preserve loop net charge while substantially altering membrane affinity, we hypothesized that local phenylalanine enrichment represents an additional sequence determinant of membrane penetration that is largely independent of electrostatic effects.

We therefore stratified all 480 simulations according to the number of local phenylalanine residues (*N_F_*) and subsequently regrouped the data within each *N_F_* category according to CBL net charge (Fig. 3E). This procedure removed the variation arising from local aromatic composition while preserving the electrostatic variation within each subgroup. For example, we regrouped the average CBL penetration depths from 210 simulations of the Phe-free C2B domains of Syt1, Syt2, Syt5, Syt6, Syt7, Syt9, and Syt10 under WT, partial charge-flip (PCF), and full charge-flip (FCF) conditions according to CBL net charge, yielding nine charge groups spanning values from −1 to 12. Syt3 C2B was excluded because it contains one local phenylalanine residue. Within each *N_F_* category, membrane penetration was strongly correlated with loop net charge, yielding very similar slope values and Pearson correlations of 0.96, 0.88, and 0.99 for *N_F_* = 0, 2, and 3, respectively (Fig. 3F). Thus, once local aromatic variation is controlled for, loop net charge alone explains the remaining variation in membrane penetration very well.

Similar charge-dependent relationships were also observed using the two independent 500 ns analysis windows (Fig. S8C-D), demonstrating that these sequence-feature relationships are robust with respect to the choice of the long-time analysis window.

### Loop net charge and local phenylalanine enrichment cooperatively govern membrane penetration across synaptotagmin C2 domains

To systematically model the two variables together, we introduced the following equation

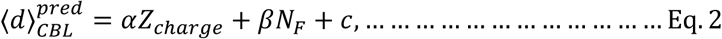

where *α*, *β*, and *c* were determined by fitting the model to the observed mean penetration depths.

The fitted coefficients (mean ± SE) were highly consistent among the three analysis time windows (*α* = 0.015 ± 0.002, 0.013 ± 0.002, 0.016 ± 0.002 nm/*Z_c_*_ℎ*arge*_ and *β* = 0.054 ± 0.006, 0.053 ± 0.006, 0.056 ± 0.007 nm/*N_F_* for the 1.5–2.5 μs, 2.0–2.5 μs, and 1.5–2.0 μs windows, respectively; Fig. 4A), demonstrating the robustness of the fitted model. The fitted coefficients indicate that an increment of *N_F_* (addition of one phenylalanine) increases penetration depth approximately as much as an increase of four charges (*Z_c_*_ℎ*arge*_ ).

**Figure 4.**
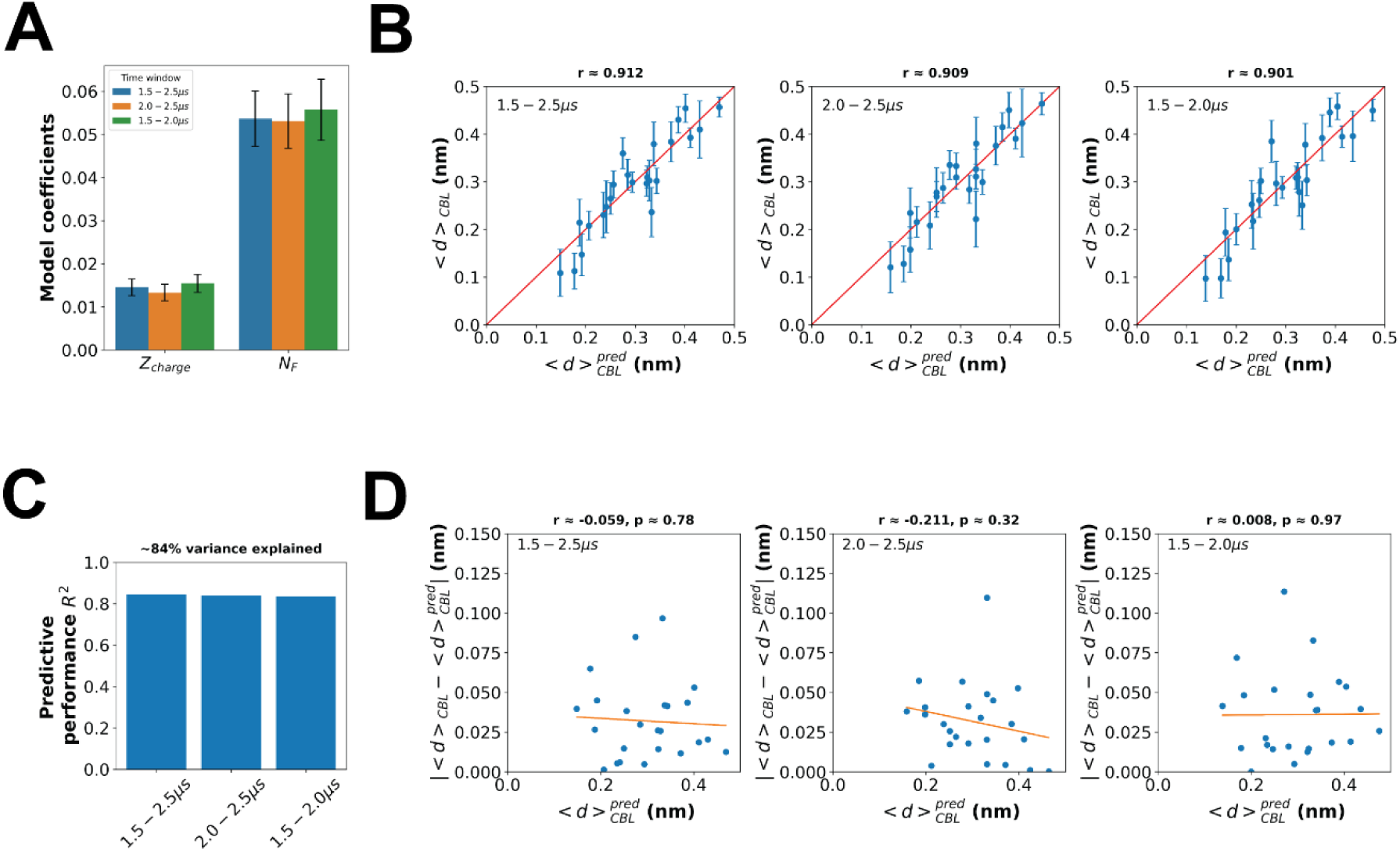
Electrostatic interactions and local aromatic residues cooperatively determine membrane penetration. **(A)** Regression coefficients associated with loop net charge (*Z_c_*_ℎ*arge*_ ) and the local number of phenylalanine residues (*N_F_*) obtained from multivariable linear regression analyses using the final 1 μs (1.5–2.5 μs, blue), the final 500 ns (2.0–2.5 μs, orange), and the preceding 500 ns (1.5–2.0 μs, green) analysis windows. Errorbars: SE. **(B)** Calibration curves comparing the observed average CBL penetration depths (⟨*d*⟩_CBL_)with those predicted by the two-feature linear model, ]inline[,where *α*, *β*, and *c* were obtained by least-squares regression. Data from the three independent analysis windows are shown. Blue dots represent the observed mean penetration depths, error bars indicate SEM, and the red line denotes perfect agreement between prediction and observation. Sample sizes for each regrouped observation are listed in Table S6. **(C)** Coefficients of determination (*R*^2^) for the two-feature regression model using the three analysis time windows, demonstrating consistent predictive performance. **(D)** Residual analysis of the two-feature model. Absolute prediction errors 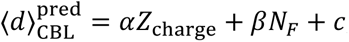 are plotted as a function of the predicted penetration depth for the three analysis windows. The residuals exhibit no systematic dependence on the predicted penetration depth (p >> 0.05), with an average absolute prediction error of approximately 0.034 nm, indicating that the remaining deviations are randomly distributed.

Incorporating local phenylalanine enrichment clearly improved the predictive performance of the model compared with the charge-only model (Fig. 3D). The predicted penetration depths were strongly correlated with the observed values, yielding Pearson correlations of approximately 0.91, 0.91, and 0.90 for the 1.5–2.5 μs, 2.0–2.5 μs, and 1.5–2.0 μs analysis windows, respectively (Fig. 4B). Accordingly, the R^2^ increased from approximately 0.59 for the charge-only model to approximately 0.84 for the two-feature model (Fig. 4C), indicating that ∼25% of the changes of penetration depth were explained by variation of *N_F_*.

Spearman correlations closely matched Pearson correlations for all three analysis windows (Fig. S9A), indicating that the predicted and observed penetration depths were related by a nearly linear monotonic relationship. Furthermore, leave-one-out validation yielded nearly identical Pearson correlations (Fig. S9B), demonstrating that the predictive performance of the model was not driven by any individual regrouped observation.

Residual analysis further showed that the absolute prediction error was independent of the predicted penetration depth across all three analysis windows (Fig. 4D), indicating that the remaining prediction errors were randomly distributed rather than systematically associated with penetration depth. Together with the small mean absolute error of approximately 0.034 nm, these results demonstrate that the two-feature model accurately captures the dominant sequence dependence of CBL membrane penetration.

Together, these results identify local phenylalanine enrichment as an independent sequence determinant of membrane penetration and establish a simple two-feature framework in which loop net charge and local phenylalanine enrichment cooperatively govern membrane penetration across synaptotagmin C2 domains.

## Discussion

### PIP₂ binding by C2 domains is primarily governed by electrostatic interactions

Experimentally, Syt1 has been shown to bind specifically to acidic membrane lipids, particularly PIP_2_ clusters at the base of SNARE proteins [19]. Ca^2+^ induces tight binding of Syt1 to membranes that involves specific interactions of the C_2_B polybasic region with PIP_2_, and it has been suggested that this disrupts Syt1-SNARE interactions [14]. It has been proposed that Syt1 interacts with PIP2 to loosely dock synaptic vesicles to the active zone in hippocampal neurons [20].

The coarse grained molecular dynamics simulation performed here, show that across all canonical Syt isoforms examined in this study, PIP₂ interactions with both CBLs and PBs were strongly associated with loop net charge. Remarkably, the charge-dependent slopes of CBL–PIP₂ and PB–PIP₂ interactions were nearly identical (∼0.28 PIP₂ molecules per unit charge (Fig. 2D)), despite the distinct locations, conformations, and biological functions of these two regions. These observations are consistent with experimental studies showing that PIP₂ binding is dominated by electrostatic interactions between negatively charged phosphoinositides and positively charged surface residues [19,21,22], making binding primarily dependent on local charge density rather than the precise structural configuration of the membrane-binding loop.

These findings provide a comprehensive physical explanation for how Ca²⁺ regulates membrane interactions of canonical Syts. Binding of Ca²⁺ to the CBLs not only neutralizes conserved acidic residues but effectively converts a locally anionic environment into a cationic one, thereby promoting the binding of polyvalent anionic phosphoinositides like PIP₂. The similar charge dependence observed for both CBLs and PBs further suggests that the primary determinant of PIP₂ binding is the net electrostatic potential generated by the protein loop rather than its precise sequence or conformation.

In contrast to PIP₂ binding, membrane penetration could not be explained by charge alone (Fig. 3C). Although increasing net charge does promote membrane insertion, charge accounted for only approximately half of the observed variation in penetration depth. This difference is physically intuitive because electrostatic attraction facilitates lipid recruitment, whereas penetration into the membrane interior additionally requires favorable hydrophobic interactions. Thus, membrane penetration is governed by both electrostatic and hydrophobic determinants, whereas PIP₂ binding is predominantly controlled by electrostatics.

### Local aromatic residues cooperate with loop net charge to govern membrane penetration

Our analyses identified local phenylalanine enrichment as a second independent sequence descriptor governing CBL membrane penetration. Incorporating this simple descriptor increased the explained change of penetration depth from approximately 57% to 82%, demonstrating that membrane penetration cannot be explained by electrostatic effects alone. Instead, loop net charge and local phenylalanine enrichment provide complementary contributions to membrane penetration across synaptotagmin C2 domains.

Sequence alignments revealed that C2A domains are frequently enriched in two to three phenylalanine residues adjacent to conserved acidic residues, whereas C2B domains predominantly contain aliphatic residues at the corresponding positions.

Because substitutions between phenylalanine and aliphatic residues generally preserve loop net charge, local phenylalanine enrichment represents an independent sequence feature that modulates membrane penetration without altering electrostatic interactions. The rigid aromatic side chain of phenylalanine is expected to stabilize hydrophobic insertion into the membrane interior, providing a qualitative physical rationale for why C2A domains exhibit stronger baseline membrane penetration than most C2B domains. This is qualitatively consistent with the White–Wimley theory that phenylalanine is ∼0.5 kcal/mol more favorable to insert into membrane then Leucine or Isoleucine [23].

The largely independent contributions of loop net charge and local phenylalanine enrichment are supported by both sequence analysis and regression modeling. Restricting the analysis to Phe-free C2B domains largely eliminated aromatic variation, allowing loop net charge alone to accurately predict membrane penetration. This is consistent with the experimental and MD simulation studies indicate that Syt7 is a membrane super-penetrator compared to Syt1 [17]. Together, these observations support a simple electrostatic–aromatic framework in which loop net charge promotes membrane engagement, whereas local phenylalanine residues stabilize membrane insertion. This two-feature framework captures the dominant sequence dependence of long-time membrane penetration and may provide a general strategy for investigating sequence-dependent membrane interactions in other C2 domains.

### Potential division of labor between canonical C2A and C2B domains in neurotransmission

Our electrostatic–aromatic framework suggests a potential functional distinction between canonical C2A and C2B domains. Because membrane penetration is jointly governed by loop net charge and local phenylalanine enrichment, whereas C2A domains are consistently enriched in phenylalanine residues adjacent to conserved acidic sites, C2A and C2B domains may contribute differently to membrane interactions during neurotransmitter release.

The phenylalanine-enriched CBLs of C2A domains may provide a relatively stable membrane anchor through hydrophobic insertion. Because this mechanism depends less strongly on charge state, C2A-mediated membrane interactions could remain partially engaged even under low-charge conditions. In contrast, C2B domains are largely depleted of local phenylalanine residues and therefore exhibit stronger dependence on charge-induced membrane interactions. Consequently, C2B domains may function as more sensitive electrostatic switches whose membrane engagement is strongly enhanced following Ca²⁺-dependent charge reversal at the CBLs.

This distinction raises the possibility that C2A and C2B domains perform complementary roles during exocytosis. C2A may help maintain membrane association through persistent hydrophobic anchoring, whereas C2B may serve as the primary electrostatic sensor that couples Ca²⁺ binding to enhanced phosphoinositide binding and membrane penetration. Such a division of labor could facilitate efficient SNARE unclamping and membrane fusion while maintaining stable membrane engagement throughout the release process.

Although this model remains speculative, it provides a simple mechanistic framework linking sequence composition, membrane interactions, and the distinct functional roles of C2A and C2B domains during neurotransmission.

### Limitations and future directions

Although the present coarse-grained simulations reveal robust properties and identify CBL net charge and local phenylalanine residues as the dominant determinants of membrane interactions, several limitations should be considered.

First, the MARTINI 2.1 force field cannot explicitly represent Ca²⁺ coordination geometry. Consequently, the partial and full charge-flip perturbations employed here were designed to systematically modulate CBL electrostatics rather than reproduce the complete structural details of Ca²⁺-bound states. Therefore, the electrostatic–aromatic framework proposed here should be viewed as a sequence-based model of membrane interactions rather than a direct description of Ca²⁺ coordination chemistry.

Second, although CBL net charge and local phenylalanine enrichment explain most of the variation in membrane penetration, substantial isoform-specific behaviors remain. For example, Syt1 and Syt5 C2B exhibited distinct responses to partial and full charge-flip perturbations despite possessing identical CBL net charges and broadly similar sequence compositions. These observations suggest that higher-order determinants, including loop flexibility, local residue packing, tandem-domain coupling, interactions beyond the CBL sequence itself, and sequence-specific conformational dynamics, may further modulate membrane interactions beyond the first-order framework identified here.

Third, the present study focused on equilibrium properties averaged over long timescales and therefore did not address transient membrane-binding kinetics or early membrane engagement processes. In addition, we occasionally observed progressive membrane burial of the β-sandwich scaffold accompanied by increased interactions between the C2 domain core and membrane headgroups. Such behavior may reflect limitations of the MARTINI 2.1 force field and should be interpreted cautiously.

Future studies using MARTINI 3 and atomistic simulations with explicit Ca²⁺ ions will be required to determine whether the electrostatic–aromatic framework identified here remains valid under realistic Ca²⁺ coordination. These simulations will also allow investigation of isoform-specific structural dynamics and higher-order sequence effects. More broadly, the simple sequence descriptors identified here provide a foundation for future machine-learning approaches aimed at uncovering sequence-dependent determinants of membrane interactions and membrane-binding kinetics. Ultimately, incorporating tandem C2AB domains, SNARE complexes, and membrane fusion intermediates may reveal how these sequence-encoded membrane interactions regulate Ca²⁺-triggered exocytosis.

## Materials and Methods

### C2B domain models

Reference structures of Syt1 (PDB: 5W5C, [15,24,25]), Syt3 (PDB: 3HN8, [26]), and Syt7 (PDB: 6ANK, [27]) were used. Structures of Syt2, Syt5, Syt6, Syt9, and Syt10 were obtained from AlphaFold models [28]. Residue ranges we used and sequence information are summarized in Supporting information.

Because explicit Ca²⁺ coordination cannot be represented in the MARTINI 2 force field, we introduced charge-flip mutations at conserved acidic residues within the Ca²⁺-binding loops (CBLs, Fig. 1A) to systematically alter CBL net charge. Partial charge-flip (PCF) variants correspond to mutation of the two conserved Ca²⁺-coordinating acidic residues (D303K/D365K in Syt1 and corresponding residues in other isoforms) (Fig. 1B) [9,17,29,30], whereas Syt3 C2B contained three substitutions (D460K/D522K/D528K) because its crystal structure shows three bound Ca²⁺ ions [26]. Full charge-flip (FCF) variants were generated by converting all five acidic residues in the CBLs to lysine.

Equivalent mutations were introduced into corresponding residues of all isoforms. Details are summarized in Fig. 1A-B. All atomistic structures were converted to MARTINI representations using martinize.py v2.5 [31].

### C2A domain models

Structures of all C2A domains were obtained from AlphaFold models (CBL sequences are shown in Fig. 1C). Residue ranges are shown in SI.

To systematically alter CBL net charge, three conserved acidic residues in Syt1 C2A (D172, D232, and D238) were converted to lysine to generate partial charge-flip (PCF, DK3) variants. Full charge-flip (FCF) variants were generated by mutating all five acidic residues in the CBLs. Equivalent mutations were introduced into corresponding residues of other isoforms (Fig. 1C-D).

### C2–Plasma Membrane (PM) System

Each C2 CBL as part of its C2 domain was inserted into an asymmetric plasma membrane by self-assembly [32] using the composition of [31,33]. After 500 steps of energy minimization, a three-stage equilibration was performed: (i) 1 ns, 0.002 ps timestep, strong x,y restraints (1000 kJ/mol/nm²); (ii) up to 20 ns, 0.02 ps timestep, weaker restraints (100 kJ/mol/nm²) with isotropic pressure coupling because of initial random orientation of lipid molecules (Some runs were terminated early to prevent full insertion of C2 into the bilayer.); and (iii) 10 ns semi-isotropic equilibration after bilayer formation. Five independent starting conditions were generated for each construct (details in https://zenodo.org/records/21284569).

### Sampling simulations

Non-inserted lipids were removed, with ions adjusted to maintain neutrality [31]. This yields an ionic strength of ∼ 110 mM. Each system was energy-minimized, equilibrated for 1 ns (0.01 ps timestep), and simulated for 2.5 µs (0.02 ps timestep). Two replicas per condition (different random seeds) were performed, yielding 10 trajectories per construct.

### Simulation details

Membrane self-assembly was run with GROMACS 2021; production simulations with GROMACS 2024 on Bridges-2 [34]. The MARTINI 2.1 force field was used. Equilibration employed a Berendsen thermostat/barostat (310 K, 1 bar, τ = 1 ps). Production runs used v-rescale thermostat (310 K, τ_T_ = 1 ps) and Parrinello–Rahman barostat (semi-isotropic, 1 bar, τ_P_ = 12 ps). Neighbor lists were updated every 20 steps with a Verlet buffer of 0.005. Simulation trajectories were recorded every 1 ns.

### Analysis

#### Syt-PIP2 binding analysis

Contacts between PIP₂ molecules and residues within the CBLs or PBs (Fig. 1E, F) were quantified every 1 ns using the minimum bead-to-bead distance. A PIP₂ molecule was considered bound when its minimum distance to any CG beads of the residue in the corresponding region was less than 0.8 nm. The total numbers of PIP₂ molecules contacting CBLs and PBs were denoted as *N*_CBL−PIP2_ and *N*_PB−PIP2_, respectively.

We characterized PIP₂ binding to CBLs and PBs by calculating average values over the last 1 μs window (1.5–2.5 μs). Additional analyses were done by calculating these values separately for the two sub time windows 1.5–2.0 μs and 2.0–2.5 μs to test the robustness of the sequence-feature analysis (see later subsections). Both intervals yielded nearly identical results, demonstrating the robustness of the reported properties.

#### CBL-PM penetration analysis

After centering the protein, a local membrane midplane was defined every 1 ns using phosphate (PO4) beads of phospholipids together with sterol (ROH) and glycolipid (GM1) reference beads located within 5 nm of the protein center. For each residue, membrane insertion depth was defined as (Eq. 1 in results section), where *Z_ref_* is the average z-position of the intracellular leaflet reference beads and *Z^min^* is the minimum z-coordinate of any beads belonging to the CBL residues. Positive values indicate penetration into the membrane interior, whereas zero or negative values indicate that the residue remains at or above the membrane headgroup region.

#### Equilibrium Quantification and Statistical Analysis

All quantities shown in bar plots were computed by averaging measurements over the final 1 μs of each simulation, to minimize the influence of long-time drift and transient fluctuations (Fig. S1). The averages calculated from the first (1.5–2.0 μs) and second (2.0–2.5 μs) halves of the final 1 μs are also provided in the Supporting Information to demonstrate the robustness of the analysis. Individual data points (black dots) represent average values from each of the n = 10 independent simulations. Generally, pairwise comparisons between individual isoforms under WT, PCF or FCF states were not performed because the objective of the study was not to identify the isoform-specific differences resulting from charge flipping, but rather to identify the dominant sequence determinants governing membrane interactions across multiple C2 domains and charge states.

#### Sequence feature analysis

Net charges of CBLs and PBs were calculated from amino acid sequences (Fig. 1A-E). To characterize local aromaticity, the number of phenylalanine (F) residues located within ±2 positions of conserved Asp and Glu residues was counted (Fig. 1A, C). These sequence-derived descriptors were used to construct SEM weighted linear regression models relating sequence features to PIP₂ binding and membrane penetration.

Pearson and Spearman correlations were calculated using all data points and performed by *pearsonr* and *spearmanr* from *scipy.stats* package. Model robustness was further assessed by leave-one-out analysis and by comparing properties obtained from the two equilibrium windows.

## Acknowledgments

This work has been supported by US National Institutes of Health (NIH) grants R35GM139608. All simulations were performed on Triton cluster from Frost Institute for Data Science and Computing (https://idsc.miami.edu/) and Bridges2 cluster from PSC and ACESS Allocation BIO250019.

## Data availability

All files to run actual simulations, code to extract data from simulation trajectories, and code to analyze data and to make figures are available in Zenodo https://zenodo.org/records/21284569. Due to their large file sizes, membrane self-assembly simulations and the trajectories generated from the production molecular dynamics simulations are not included in this repository. These files are available from the corresponding author(s) upon reasonable request.

## Declaration of generative AI and AI-assisted technologies in the writing process

During the preparation of this work, the first author used ChatGPT to improve language and readability. After using this tool/service, the author(s) reviewed and edited the content as needed and took full responsibility for the content of the publication.

