## Supplemental information for "Electrostatics and Local Aromatic Residues Govern Lipid Binding and Membrane Penetration of Synaptotagmin C2 Domains"

### Supporting information

#### Supporting Methods

Reference structures of Syt1 residues 270–418, Syt3 residues 427-573, and Syt7 residues 266-403 from crystal structures were used. Structures of Syt2 (Uniport: P29101), Syt5 (Uniport: P47861), Syt6 (Uniport: Q62746), Syt9 (Uniport: Q925C0) and Syt10 (Uniport: O08625) were generated by AlphaFold and residues 271-419 of Syt2, residues 237-373 of Syt5, residues 360-498 of Syt6, residues 350-491 of Syt9, and residues 361-499 of Syt10 were used, respectively.

All C2A models for Syt1 (residues 141-261), Syt2 (142-262), Syt3 (299-419), Syt5 (108-228), Syt6 (230-352), Syt7 (136-256), Syt9 (222-342) and Syt10 (231-353) are taken from Alphafold generated model (Fig. 1C).

All simulations were conducted for 2.5 μs. Following production runs, trajectories were preprocessed prior to analysis. Specifically, each trajectory was centered on the C2 domain to remove translational drift under periodic boundary conditions. Solvent molecules and ions were removed to facilitate protein–lipid interaction analysis and reduce computational overhead during data extraction.

Trajectory processing and quantitative analyses were performed using *MDTraj* (v1.10.0). All downstream data analysis and figure generation were carried out in Python using the following packages: *NumPy* (v1.24.4), *Pandas* (v1.5.3), *Matplotlib* (v3.7.5), *Seaborn* (v0.11.2), and *Statannotations* (v0.6.0).

Further methodological details, including analysis scripts and workflow documentation, are available at: DYAD_LINK.

#### Supporting Table Legends

**Table S1.** Individual CBL–$PIP_2$ binding values from independent simulations.

Individual CBL–$PIP_2$ binding values averaged over the three analysis windows (1.5–2.5 μs, 2.0–2.5 μs, and 1.5–2.0 μs) for all independent simulations.

**Table S2.** Individual PB–$PIP_2$ binding values from independent simulations.

Individual PB–$PIP_2$ binding values averaged over the three analysis windows (1.5–2.5 μs, 2.0–2.5 μs, and 1.5–2.0 μs) for all independent simulations.

**Table S3.** Sample sizes and statistics for CBL–$PIP_2$ binding grouped by loop net charge.

Numbers of regrouped observations in each loop net charge ($Z_{\text{charge}}$) category used for the regression analyses of CBL–$PIP_2$ binding. Mean ± SEM values over the three analysis windows (1.5–2.5 μs, 2.0–2.5 μs, and 1.5–2.0 μs) were also included.

**Table S4.** Sample sizes and statistics for PB–$PIP_2$ binding grouped by PB net charge.

Numbers of regrouped observations in each PB net charge ($Z_{\text{charge}}$) category used for the regression analyses of PB–$PIP_2$ binding. Mean ± SEM values over the three analysis windows (1.5–2.5 μs, 2.0–2.5 μs, and 1.5–2.0 μs) were also included.

**Table S5.** Individual membrane penetration depths.

Individual CBL membrane penetration depths averaged over the three analysis windows (1.5–2.5 μs, 2.0–2.5 μs, and 1.5–2.0 μs) for all independent simulations.

**Table S6.** Sample sizes and statistics for the two-feature membrane penetration model.

Numbers of regrouped observations in each combination of loop net charge ($Z_{\text{charge}}$) and local phenylalanine number ($N_F$) used for the multivariable regression analyses of membrane penetration. Mean ± SEM values over the three analysis windows (1.5–2.5 μs, 2.0–2.5 μs, and 1.5–2.0 μs) were also included.

#### Supporting Figures

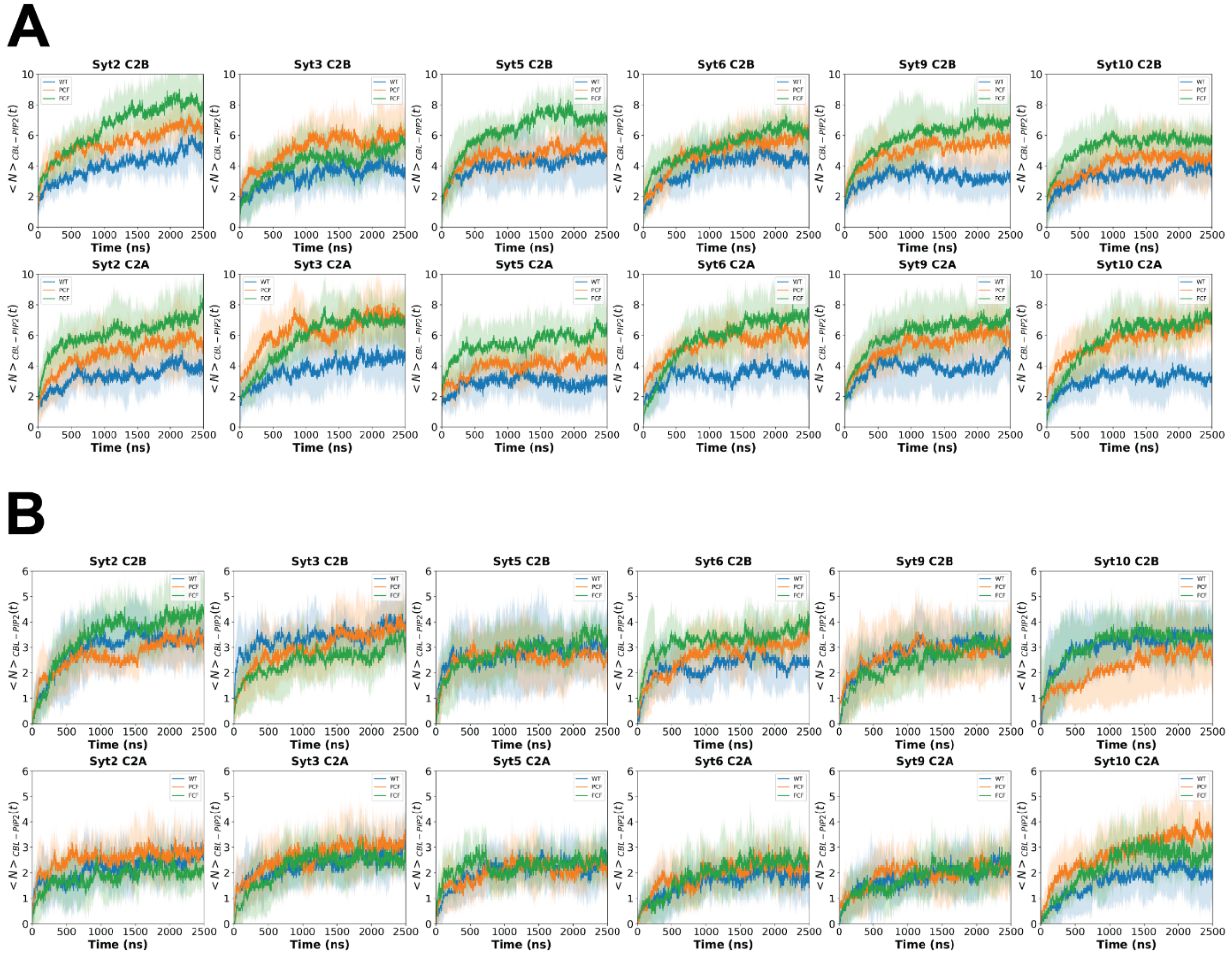


**Figure S1. Representative long-time trajectories of $PIP_2$ binding.**

**(A)** Long-time trajectories of CBL–$PIP_2$ binding for the remaining Syt isoforms (Syt2, Syt3, Syt5, Syt6, Syt9, and Syt10) under WT, PCF, and FCF conditions. Solid lines represent the mean values from $n$ = 10 independent simulations, and shaded regions indicate standard deviations.

**(B)** Long-time trajectories of PB–$PIP_2$ binding for the same simulations. Solid lines represent the mean values from $n$ = 10 independent simulations, and shaded regions indicate standard deviations.

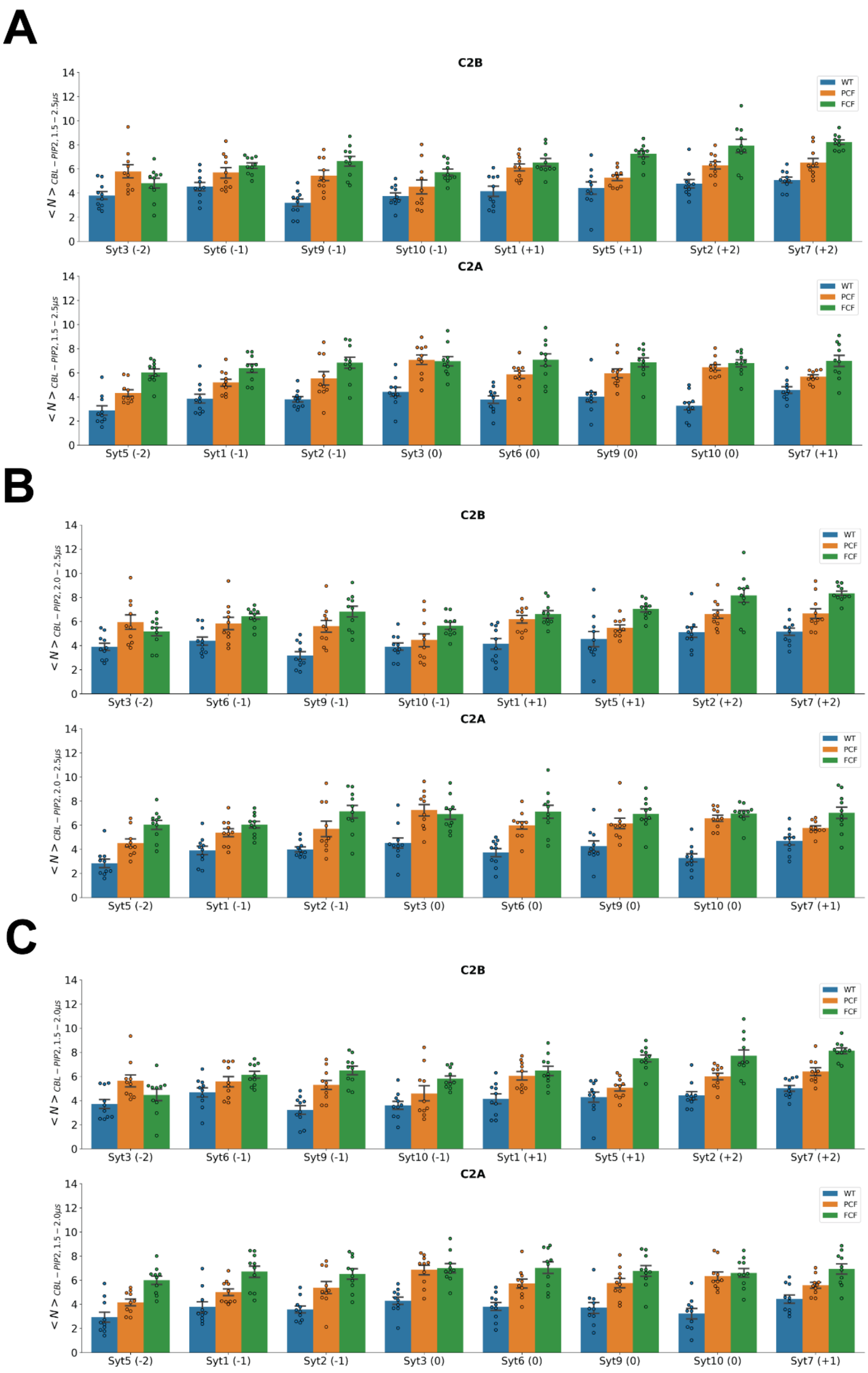
A
C2B
C2A
B
C2B
C2A
C
C2B
C2A
WT
PCF
FCF
< N > CBL − PIP2, 1.5 − 2.5μs
< N > CBL − PIP2, 2.0 − 2.5μs
< N > CBL − PIP2, 1.5 − 2.0μs
Syt3 (-2)
Syt6 (-1)
Syt9 (-1)
Syt10 (-1)
Syt1 (+1)
Syt5 (+1)
Syt2 (+2)
Syt7 (+2)
Syt5 (-2)
Syt1 (-1)
Syt2 (-1)
Syt3 (0)
Syt6 (0)
Syt9 (0)
Syt10 (0)
Syt7 (+1)

**Figure S2.** PIP2 binding of CBLs across synaptotagmin isoforms.

**(A-C)** Average numbers of PIP2 molecules contacting CBLs during **(A)** 1.5–2.5 μs, **(B)** 2.0–2.5 μs and **(C)** 1.5–2.0 μs.

Bars represent mean ± SEM and dots correspond to ten independent simulations. Numbers in parentheses indicate net charges of CBLs or PBs. All individual PIP2 binding are summarized in Table S1.

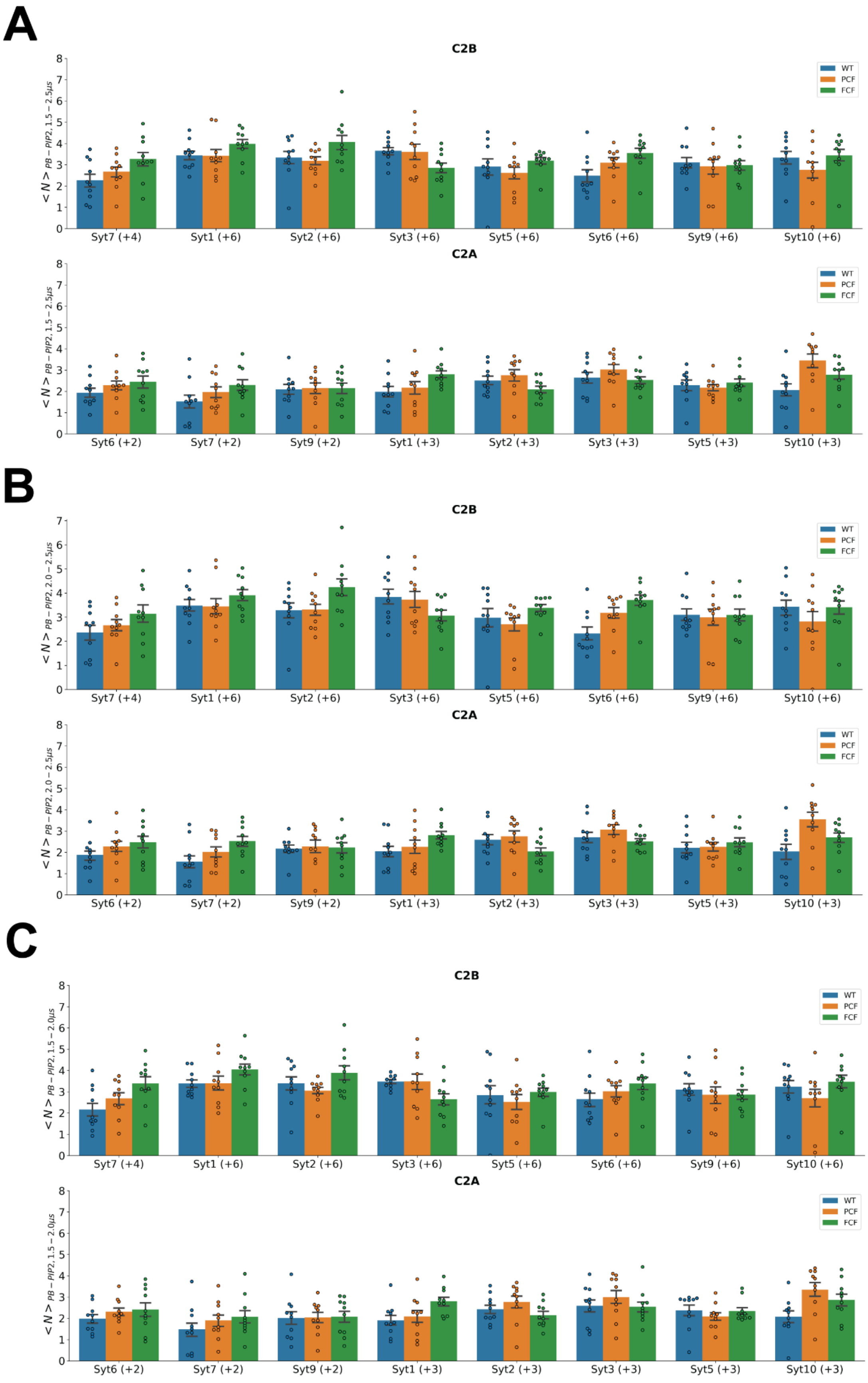
A
C2B
WT
PCF
FCF
< N > PB − PIP2, 1.5 − 2.5μs
Syt7 (+4)
Syt1 (+6)
Syt2 (+6)
Syt3 (+6)
Syt5 (+6)
Syt6 (+6)
Syt9 (+6)
Syt10 (+6)
C2A
Syt6 (+2)
Syt7 (+2)
Syt9 (+2)
Syt1 (+3)
Syt2 (+3)
Syt3 (+3)
Syt5 (+3)
Syt10 (+3)
B
< N > PB − PIP2, 2.0 − 2.5μs
C
< N > PB − PIP2, 1.5 − 2.0μs

**Figure S3.** PIP2 binding of CBLs across synaptotagmin isoforms.

**(A-C)** Average numbers of PIP2 molecules contacting PBs during **(A)** 1.5–2.5 μs, **(B)** 2.0–2.5 μs and **(C)** 1.5–2.0 μs.

Bars represent mean ± SEM and dots correspond to ten independent simulations. Numbers in parentheses indicate net charges of CBLs or PBs. All individual PIP2 binding are summarized in Table S2.

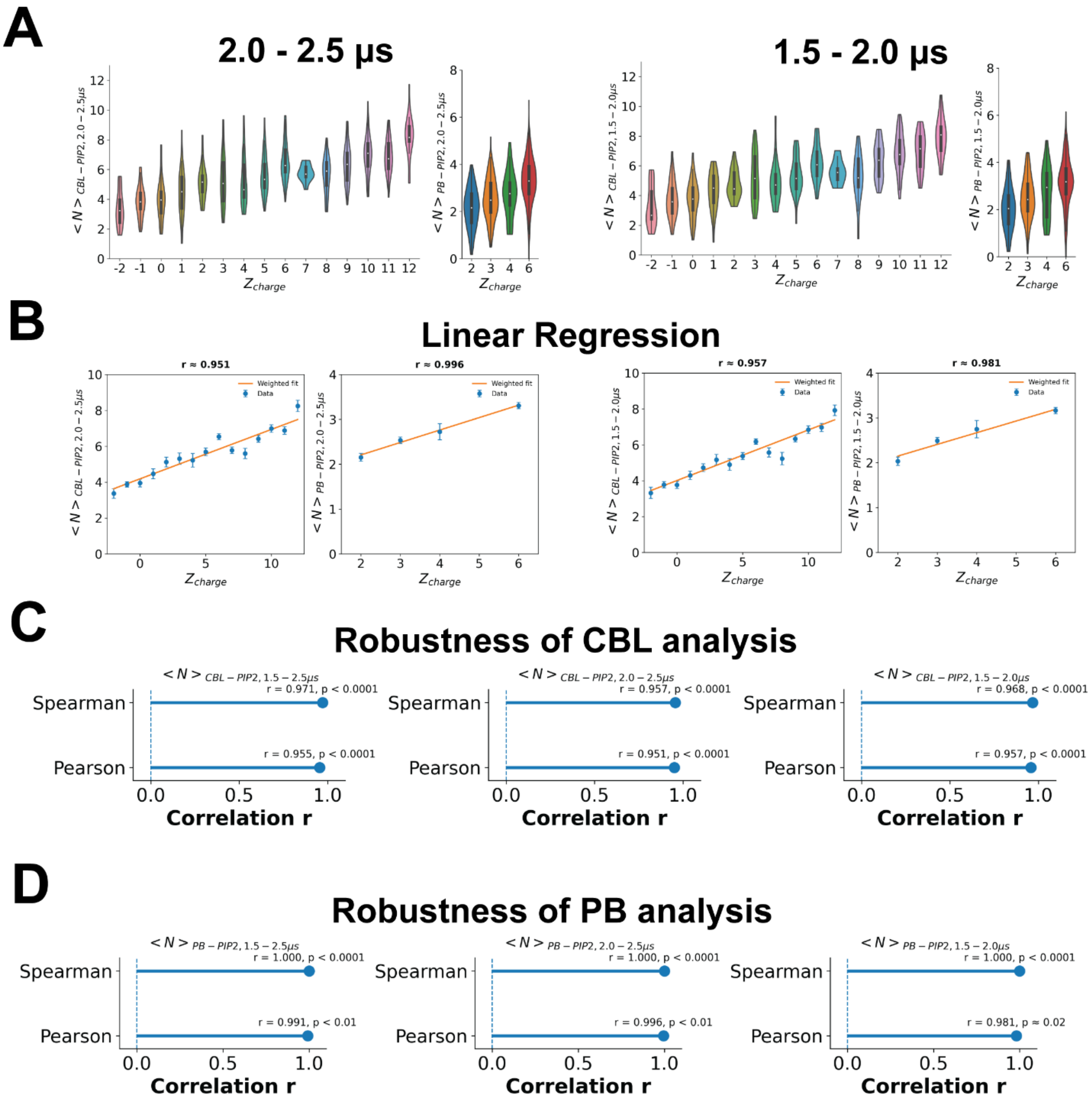


**Figure S4. Robustness of the sequence-feature analysis with respect to the choice of the analysis window.**

**(A)** Distributions of the average number of $PIP_2$ molecules bound to calcium-binding loops (CBLs, left) and polybasic patches (PBs, right) during the final 500 ns (2.0–2.5 μs) and the preceding 500 ns (1.5–2.0 μs). Data from all C2A and C2B domains, Syt isoforms, and charge states (WT, PCF, and FCF) were regrouped according to the corresponding net charge ($Z_{charge}$).

**(B)** Linear regression analyses between the mean number of bound $PIP_2$ molecules and loop net charge using the two independent 500 ns analysis windows. Blue dots represent the mean values averaged across all C2 domains sharing the same net charge, and error bars indicate SEM. Orange lines indicate the best-fit linear regressions.

**(C)** Pearson and Spearman correlation coefficients describing the relationship between CBL $PIP_2$ binding and loop net charge for the final 1 μs (1.5–2.5 μs), the final 500 ns (2.0–2.5 μs), and the preceding 500 ns (1.5–2.0 μs).

**(D)** Pearson and Spearman correlation coefficients describing the relationship between PB $PIP_2$ binding and PB net charge for the same three analysis windows.

The nearly identical distributions, regression relationships, and correlation coefficients obtained from the three independent analysis windows demonstrate that the sequence-feature analysis is robust with respect to the choice of the  analysis window.

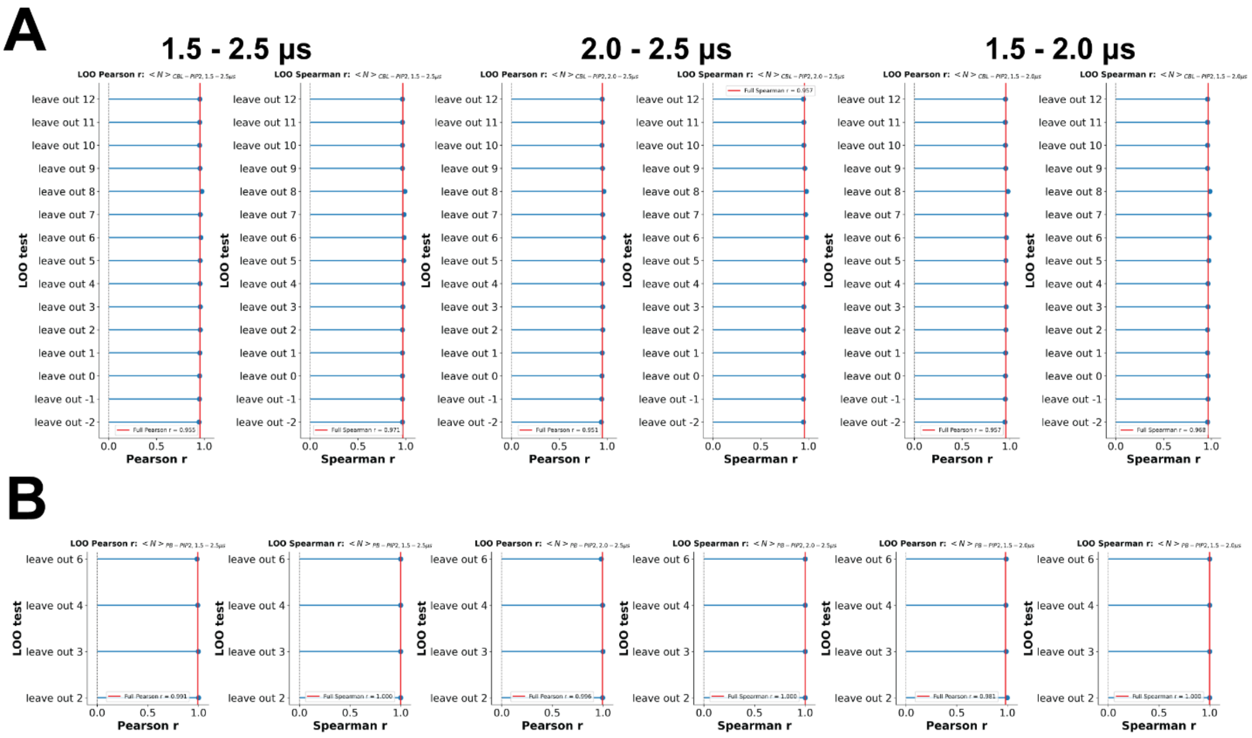


**Figure S5. Leave-one-out validation demonstrates the robustness of the sequence-feature correlations.**

**(A)** Leave-one-out (LOO) Pearson and Spearman correlations between CBL–$PIP_2$ binding and CBL net charge. In each LOO analysis, one net-charge group was excluded before recalculating the correlation coefficients.

**(B)** Leave-one-out (LOO) Pearson and Spearman correlations between PB–$PIP_2$ binding and PB net charge. In each LOO analysis, one net-charge group was excluded before recalculating the correlation coefficients.

Red vertical lines indicate the correlation coefficients obtained using the complete dataset.

Together with Fig. S4, these analyses demonstrate that the reported sequence-feature relationships are robust with respect to both the choice of the analysis window and the inclusion of individual charge groups.

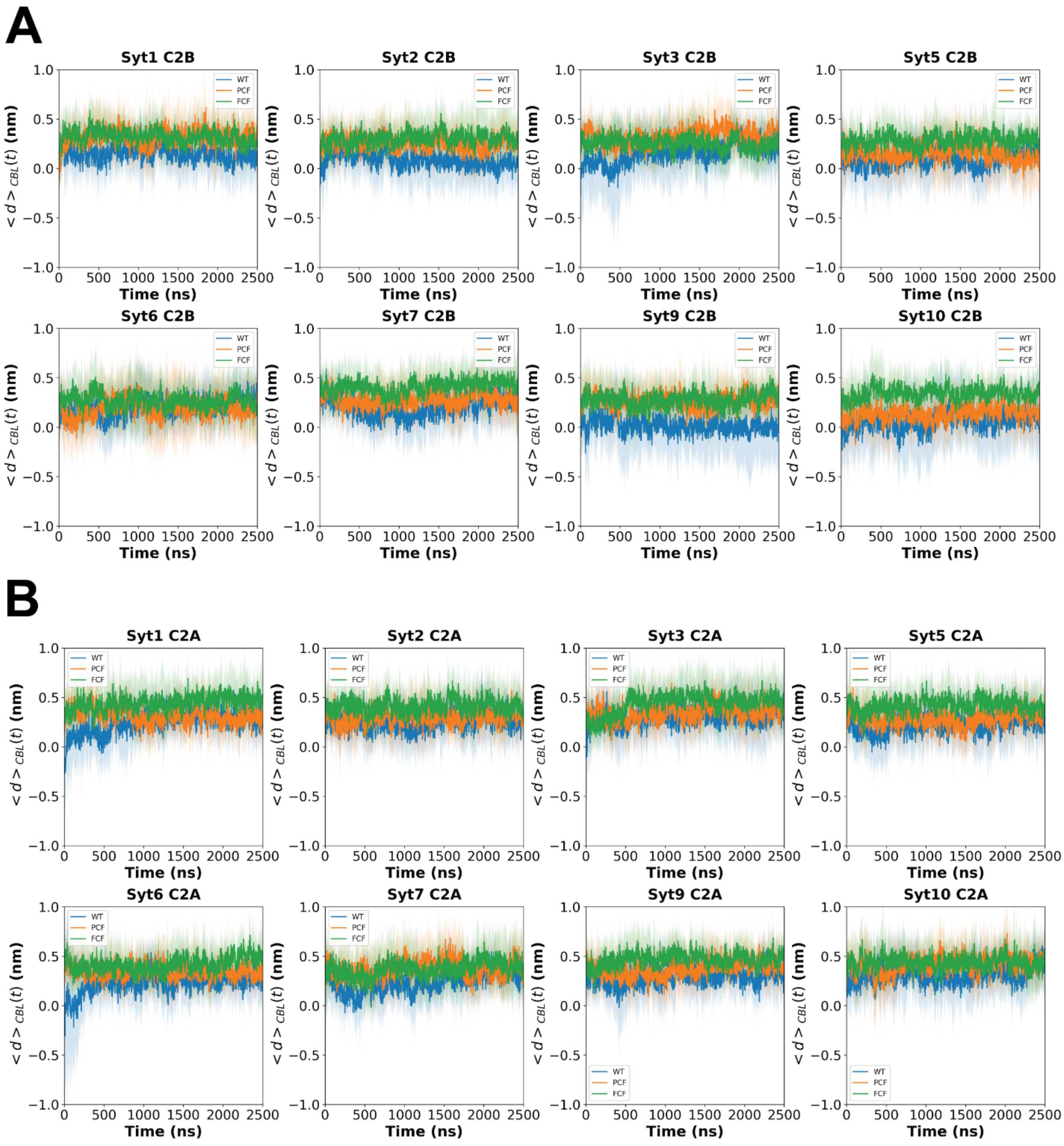


**Figure S6. Long-time trajectories of CBL membrane penetration for all synaptotagmin isoforms.**

**(A)** Mean CBL membrane penetration trajectories of the C2B domains under WT, PCF, and FCF conditions.

**(B)** Mean CBL membrane penetration trajectories of the C2A domains under WT, PCF, and FCF conditions.

Solid lines represent the mean values from $n$ = 10 independent simulations, and shaded regions indicate standard deviations.

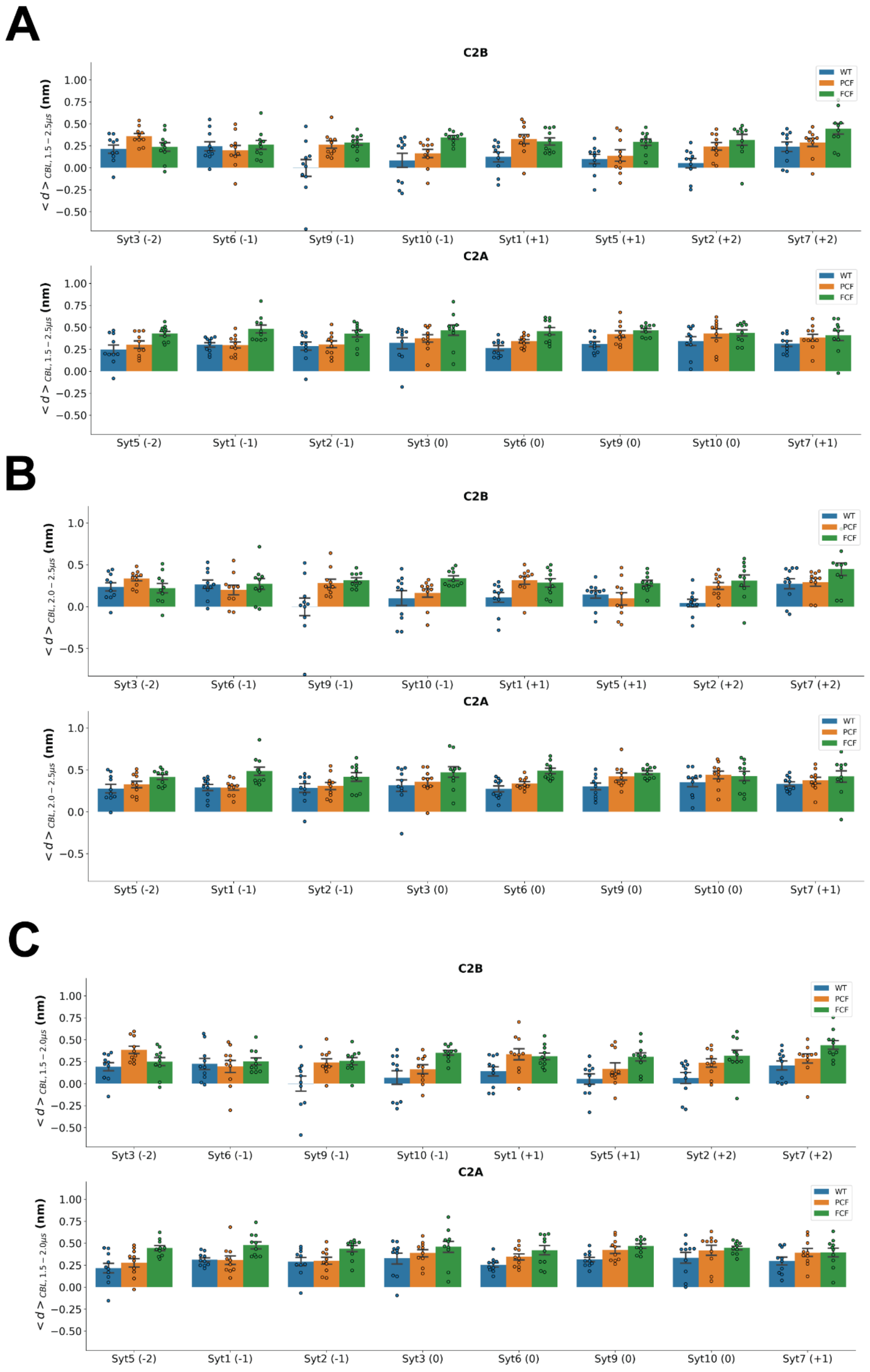

A
C2B
WT
PCF
FCF
$< d >_{CBL,\ 1.5-2.5 \mu s}$ (nm)
1.00
0.75
0.50
0.25
0.00
−0.25
−0.50
Syt3 (-2)
Syt6 (-1)
Syt9 (-1)
Syt10 (-1)
Syt1 (+1)
Syt5 (+1)
Syt2 (+2)
Syt7 (+2)
C2A
Syt5 (-2)
Syt1 (-1)
Syt2 (-1)
Syt3 (0)
Syt6 (0)
Syt9 (0)
Syt10 (0)
Syt7 (+1)
B
$< d >_{CBL,\ 2.0-2.5 \mu s}$ (nm)
1.0
0.5
0.0
−0.5
C
$< d >_{CBL,\ 1.5-2.0 \mu s}$ (nm)

**Figure S7. CBL penetration depths of all Syt isoforms.**

**(A-C)** Average numbers of CBL penetration depths during **(A)** 1.5–2.5 μs, **(B)** 2.0–2.5 μs and **(C)** 1.5–2.0 μs.

Bars represent mean ± SEM and dots correspond to ten independent simulations. Numbers in parentheses indicate net charges of CBLs or PBs. All individual CBL penetration depths are summarized in Table S5.

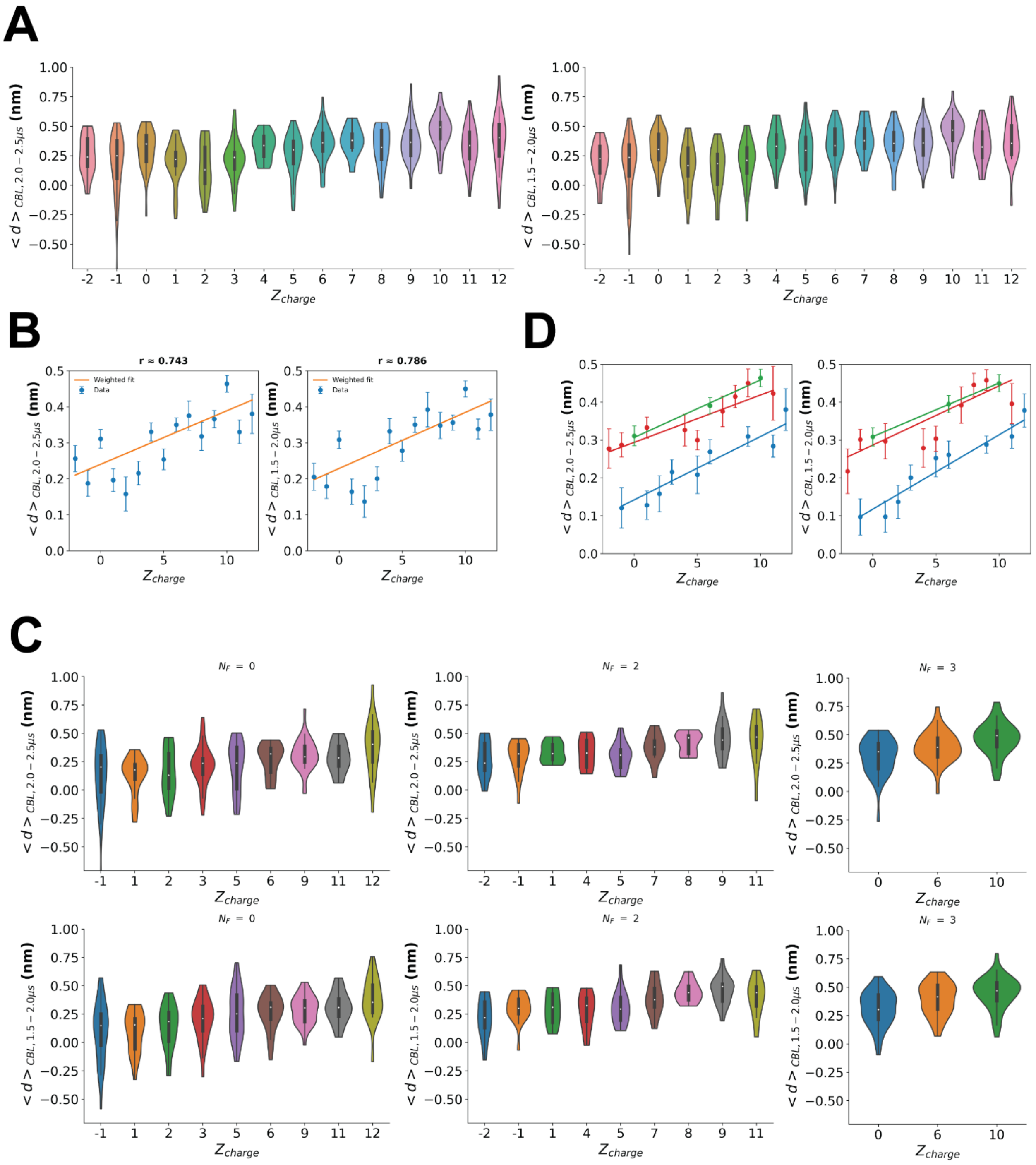

**Figure S8. Robustness of the charge- and phenylalanine-dependent membrane penetration analysis with respect to the choice of the analysis window**.

**(A)** Distributions of the average membrane penetration depth during the final 500 ns (2.0–2.5 μs, left) and the preceding 500 ns (1.5–2.0 μs, right). Data from all C2A and C2B domains, Syt isoforms, and charge states (WT, PCF, and FCF) were regrouped according to the CBL net charge ($Z_{charge}$).

(B) Linear regressions between the mean membrane penetration depth and CBL net charge using the two independent 500 ns analysis windows. Blue dots represent the mean penetration depths averaged across all C2 domains sharing the same net charge, and error bars indicate SEM. Orange lines indicate the best-fit linear regressions.

(C) Distributions of membrane penetration depth after regrouping according to both CBL net charge ($Z_{charge}$) and the local phenylalanine enrichment ($N_F$) for the final 500 ns (top) and the preceding 500 ns (bottom). The same charge-dependent trends within each $N_F$ category are observed in both analysis windows.

(D) Linear regressions between membrane penetration depth and CBL net charge for the three $N_F$ categories ($N_F$ = 0, 2, and 3) using the two independent 500 ns analysis windows. The slopes are 0.017 ± 0.003, 0.013 ± 0.003, and 0.015 ± 0.002 nm/$Z_{charge}$ for $N_F$ = 0, 2, and 3 for the final 500 ns (left), and the slopes are 0.020 ± 0.002, 0.016 ± 0.004, and 0.014 ± 0.0002 nm/$Z_{charge}$ for $N_F$ = 0, 2, and 3 for the preceding 500 ns (bottom).

The nearly identical charge-dependent distributions, regression relationships, and $N_F$-dependent trends obtained from the two independent analysis windows demonstrate that the two-feature (loop net charge and local phenylalanine enrichment) framework is robust with respect to the choice of the analysis window.

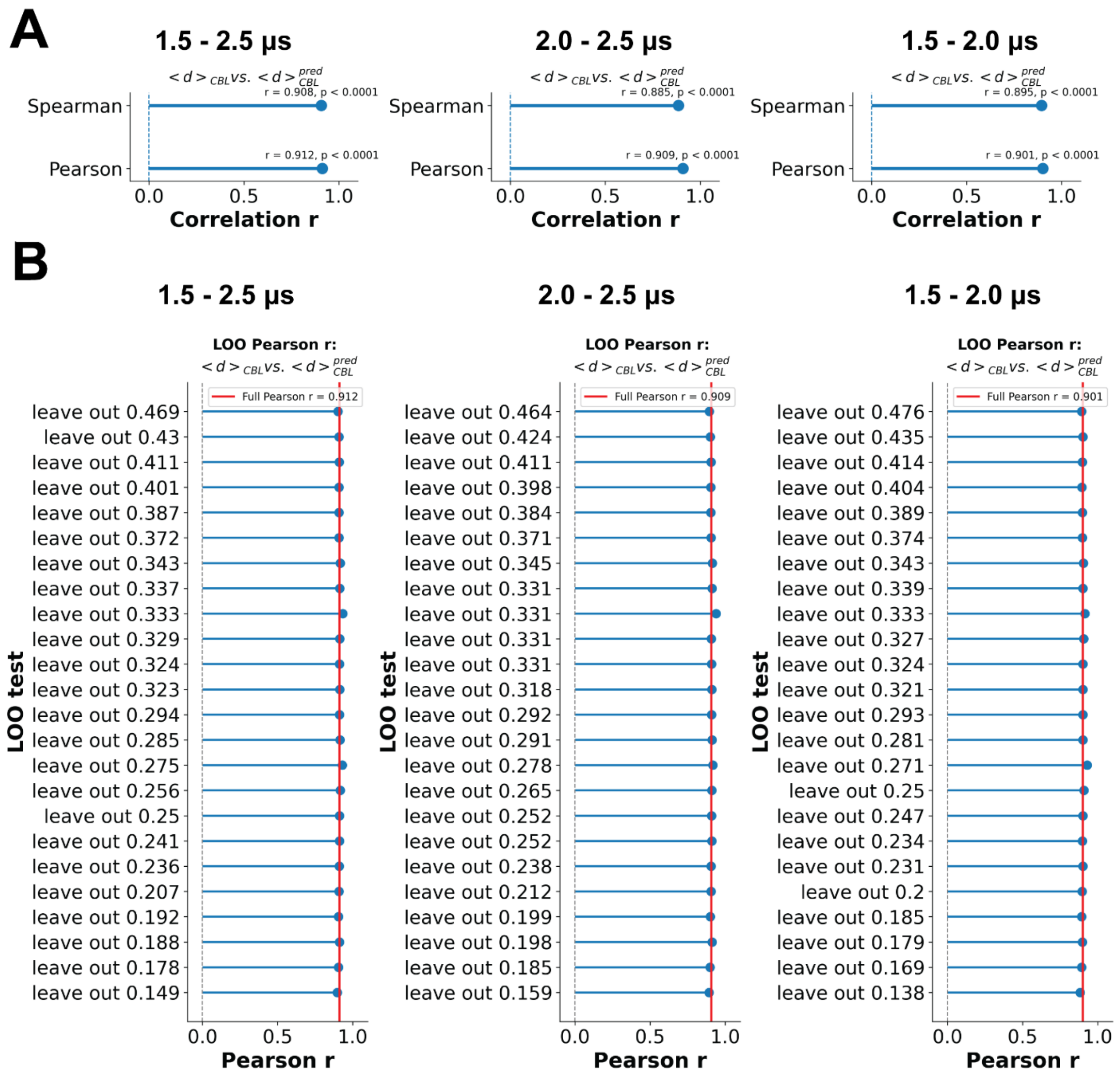


**Figure S9.** Validation of the two-feature model for predicting membrane penetration.

**(A)** Pearson and Spearman correlations between the observed membrane penetration depths and those predicted by the two-feature model (loop net charge and local phenylalanine enrichment, $N_F$) using the final 1 μs (1.5–2.5 μs), the final 500 ns (2.0–2.5 μs), and the preceding 500 ns (1.5–2.0 μs) analysis windows.

**(B)** Leave-one-out (LOO) validation of the two-feature prediction model using the three analysis windows. In each LOO analysis, one regrouped observation (corresponding to a specific combination of loop net charge and local phenylalanine enrichment) was excluded before refitting the linear model and recalculating the Pearson correlation coefficient. Red vertical lines indicate the Pearson correlation coefficients obtained using the complete datasets.

The consistently high prediction accuracies and the minimal changes observed during leave-one-out validation demonstrate that the predictive performance of the two-feature model is robust with respect to both the choice of the analysis window and the inclusion of individual regrouped observations.
